# The catalytic rate constant of *Trichoderma longibrachiatum* family 7 cellobiohydrolase on cellulose

**DOI:** 10.1101/2023.08.09.552603

**Authors:** Tina Jeoh

**Affiliations:** Department of Biological and Agricultural Engineering, University of California Davis, One Shields Avenue, Davis, CA 95616 United States

**Keywords:** Cellobiohydrolases, catalytic rate constant, *Trichoderma longibrachiatum* CBHI, Cel7, k_cat_

## Abstract

Cellobiohydrolases are considered ‘workhorse’ cellulases in the saccharification of cellulosic substrates. As solubilized cellobiose products are easily measured, cellobiohydrolases have been useful tools in studies of cellulose hydrolysis. Access to purified cellobiohydrolases with known catalytic rate constant (k_cat_), or the turnover rate, is particularly useful. Here, we determined the k_cat_ of the commercially available *Trichoderma longibrachiatum* CBHI at 30^°^C, 40^°^C, and 50^°^C to be 0.5 s^-1^, and 2.6 s^-1^, and 5.3 s^-1^, respectively. The energy of activation of productively bound *Tl*CBHI was estimated to be 93 ± 17 kJ/mol.

## Introduction

*Trichoderma longibrachiatum* CBHI (*Tl*CBHI, EC 3.2.1.176) is a reducing-end specific, glycoside hydrolase family 7 (GH7) cellobiohydrolase (www.cazy.org)^1^. *Tl*CBHI is commercially available as a monocomponent, i.e. purified form; as such, *Tl*CBHI is useful in bioassays studying enzymatic mechanisms of cellulose hydrolysis. *Tl*CBHI is a similar enzyme to *Trichoderma reesei* Cel7A (*Tr*Cel7A) in that both target cellulose from the reducing ends, and hydrolyze glycosidic bonds to release soluble cellobiose. While the intrinsic catalytic rate constant, k_cat_ of *Tr*Cel7A has been shown to be ∼ 5 s^-1^ by experimental and computational determinations^2–4^, the k_cat_ of *Tl*CBHI on cellulose remains unmeasured.

Productive binding sites on cellulose are those at which cellulase enzymes can form an active complex with cellulose, hydrolyze glycosidic bonds, thereby generating product ^5^. For reducing-end specific cellobiohydrolases, productive binding sites are accessible reducing ends where a complexed enzyme can release soluble cellobiose. Previous work has shown that the initial availability of accessible productive Cel7A binding sites on cellulose varies with the source and upstream treatment ^6^. Moreover, that the catalytic domain of Cel7A (Cel7ACD) access fewer productive binding sites on microcrystalline cellulose than fully intact Cel7A equipped with its cellulose binding module and linker ^7^. Hydrolysis rates of cellulose depends on how successful cellulases are complexing to productive binding sites on cellulose. Thus, both substrate properties, i.e. availability of productive binding sites, and enzyme properties, i.e. ability to complex to potential productive binding sites, can limit cellulose hydrolysis rates.

The rate at which productively bound cellobiohydrolase hydrolyzes cellulose is first order with respect to the rate at which cellobiose is released into solution ^8^. The initial rate of cellobiose release can be attributed to the ‘first cut’ made by productively complexed cellobiohydrolases. Thus, the initial rate of cellobiose release when all productive binding sites are occupied, i.e. the maximum initial cellobiose release rate when the substrate is saturated, informs on the productive cellulase binding capacity of the cellulosic substrate. Previously, we established a method to determine the productive binding capacity of cellulosic substrates from initial cellobiose release rates catalyzed by TrCel7A with known k_cat_ at saturation, measured using an amperometric cellobiose biosensor ^5,6^.

Here, we determined the k_cat_ of *Tl*CBHI by measuring initial cellobiose release rates on swollen cellulose of known productive binding capacity under saturating conditions. We also estimated the energy of activation of cellulose hydrolysis by *Tl*CBHI from k_cat_ estimates at various temperatures.

## Methods

### Enzymes

*Trichoderma longibrachiatum* CBHI (*Tl*CBHI) was purchased from Neogen (E-CBHI, Neogen Corporation, Lansing, MI) and buffer exchanged into 50 mM sodium acetate, pH 5.0 prior to use. Stock enzyme concentration was measured by absorbance at 280 nm and calculated using enzyme molecular weight of 54,319.34 Da and molar extinction coefficient, ε280 = 11960 M^-1^ cm^-1^. Enzyme stocks were stored refrigerated.

*Trichoderma reesei* Cel7A (*Tr*Cel7A) was purified from a commercial preparation (Sigma Cat# C2730) by anion exchange, affinity capture on a *p-*aminophenyl cellobioside (*p*APC) column, and size exclusion as previously described ^9^.

### Swollen cellulose

Swollen cellulose was regenerated from swelling filter paper (Whatman #1) in concentrated phosphoric acid ^10^. Filter paper (5 g), cut into small pieces, was stirred continuously in 250 mL of cold 85% phosphoric acid at 4 ^°^C overnight. The phosphoric acid swollen cellulose (PASC) was precipitated by pouring into a large volume of ice-cold water. The swollen cellulose was washed repeatedly with water until near neutral pH, suspended in water and autoclaved. The concentration of swollen cellulose was determined by the Anthrone assay against a glucose standard curve ^11^.

### Hydrolysis time course of swollen cellulose by TlCBHI and TrCel7A

Swollen cellulose was hydrolyzed over the course of 24 hours by *Tl*CBHI and *Tr*Cel7A at a loading of 10 μmoles/g with 0.15 mg/mL cellulose in 50 mM sodium acetate buffer, pH 5.0, at 50^°^C with end-over-end mixing. Reactions were stopped by filtering with a 0.45 μm filter plate (Pall Life Sciences, PN 8048). Reaction progress was determined by measuring reducing sugars using the bicinchoninic colorimetric assay (BCA) against a cellobiose standard curve ^12^. Four replicates were conducted of each reaction.

### Determining saturating concentrations of productively bound TlCBHI on swollen cellulose

Saturating concentrations of productively bound *Tl*CBHI on swollen cellulose was determined from maximum cellobiose release rates under saturating conditions as previously described ^6^. Initial (maximum) cellobiose release at varying enzyme/cellulose loading were determined from time-resolved cellobiose measurements made with a cellobiose dehydrogenase amperometric biosensor. Each reaction, at enzyme/substrate loadings in the range of 40 – 350 μmole/g cellulose, contained 0.05 mg/mL swollen cellulose in 50 mM sodium acetate buffer (pH 5.0) in 1.5 mL reaction volume. Substrate and buffer were equilibrated in the jacketed stir cell with the biosensor probe at predetermined reaction temperatures, and the reaction was initiated by the addition of the *Tl*CBHI stock to the target enzyme/substrate loading. Amperometric signal was recorded at a 10 per second sampling rate for up to ∼30 seconds and converted to cellobiose concentration via standard curves measured separately. The steepest, initial slope of the cellobiose v. time curve determined the initial/maximum rate of cellobiose release (r_CB_). Saturating concentrations of productively bound *Tl*CBHI were determined empirically from fits to the following relationship using the curve fitting toolbox in Matlab (Mathworks, Inc., Natick, MA):

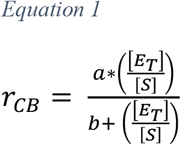

where, r_CB_ = rate of cellobiose release (μM/s), [E_T_]/[S] = enzyme loading (μmole/g), and a and b are fitting parameters, corresponding to the estimate of the maximum cellobiose release rate when all productive binding sites are occupied (r_CB_|_max_), and the half-saturation constant, respectively. Measurements were conducted at 30, 40 and 50^°^C.

### Estimating the catalytic rate constant, k_cat_ of TlCBHI

The rate of cellobiose release is directly proportional to and first order with respect to the concentration of productively bound enzyme ([E_BP_]), and the saturation concentration of productively bound enzymes was previously defined to be the productive binding capacity of the cellulosic substrate ^5^. Thus, the k_cat_ of *Tl*CBHI can be estimated as follows:

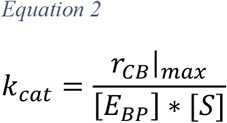

where, k_cat_ = the catalytic rate constant (s^-1^), [E_BP_] = concentration of productively bound enzyme (μmole/g), and [S] = initial cellulose concentration (g/L).

### Estimating the energy of activation of cellulose hydrolysis by TlCBHI

The energy of activation, E_a_, of cellulose hydrolysis by productively bound *Tl*CBHI was estimated from the slope of the natural logarithm of the Arrhenius equation:

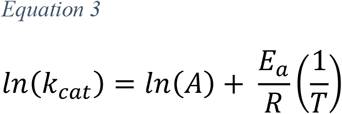

Where A = the preexponential factor, E_a_ = the energy of activation (kJ/mol), R = universal gas constant (8.3145 × 10^−3^ kJ/mol/K), and T = temperature (K).

## Results and Discussion

### The productive TlCBHI binding capacity of swollen cellulose

The productive *Tr*Cel7A binding capacity of swollen cellulose was previously determined to be 5.75 ± 0.45 μmoles/g ^5,6^. An enzyme loading of 10 μmoles/g in a hydrolysis reaction is thus considered ‘substrate-limiting’ where the initial availability of enzymes exceeds available productive binding sites by > 1.7 fold. At this loading, *Tr*Cel7A fully hydrolyzed the swollen cellulose by 16 hours (Figure 1). *Tl*CBHI hydrolysis of the swollen cellulose tracked that of *Tr*Cel7A until at least 4 hours (34 ± 0.6% and 36 ± 0.3% conversion, respectively), beyond which, hydrolysis by *Tl*CBHI slowed down to reach a maximum of 78 ± 2.1% at 24 hours. Similarity in swollen cellulose hydrolysis rates by the two enzymes suggests that *Tl*CBHI and *Tr*Cel7A found similar access to productive binding sites early on. The hydrolysis slowdown by *Tl*CBHI suggests that as the reaction proceeds, there were potential productive binding sites that were accessible to *Tr*Cel7A but not *Tl*CBHI. The hydrolysis slowdown is not likely due to product inhibition at the low soluble sugar concentrations (up to ∼0.5 mM), but it is possible that *Tl*CBHI becomes inactivated sooner than *Tr*Cel7A.

**Figure 1:**
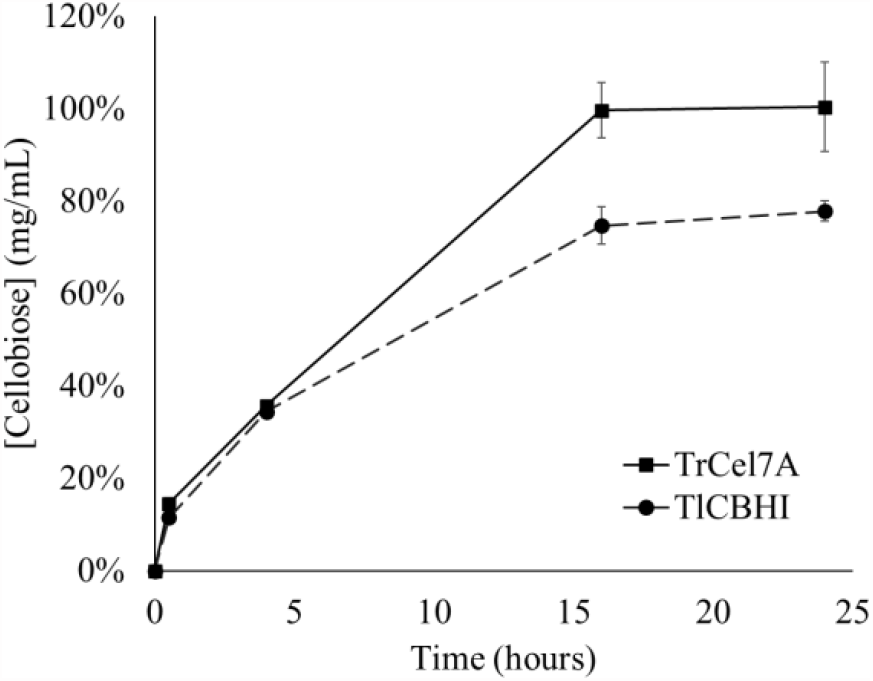
Hydrolysis time course of swollen cellulose by 10 μmoles/g of TrCel7A and TlCBHI in 50 mM sodium acetate buffer, pH 5, at 50^°^C with end-over-end rotation.

Considering that under the substrate-limited reaction condition, *Tl*CBHI exhibited similar initial swollen cellulose hydrolysis as *Tr*Cel7A, we can assume that the initial productive *Tl*CBHI binding capacity of swollen cellulose is also 5.75 μmoles/g.

### The catalytic rate constant of Trichoderma longibrachiatum CBHI (TlCBHI)

Initial cellobiose release rates from *Tl*CBHI hydrolysis of cellulose were determined from within the first ∼ 5 seconds of the reaction at various enzyme loadings at 30, 40 and 50^°^C (Figure 2). Maximum cellobiose release rates, i.e. d[CB]/dt where all productive binding sites were saturated, were estimated by fitting an empirical saturation equation to estimate the parameter *a* (Equation 1, Figure 2). Using a productive *Tl*CBHI binding capacity of swollen cellulose of 5.75 μmoles/g ^5,6^, the estimated the catalytic rate constant, k_cat_ of *Tl*CBHI at 30 ^°^C, 40 ^°^C, and 50 ^°^C were 0.5 s^-1^, and 2.6 s^-1^, and 5.3 s^-1^, respectively. Relating the k_cat_ values with temperature via the Arrhenius relationship (Equation 3) estimated the energy of activation (E_a_) of 93 ± 17 kJ/mol, within the order of magnitude of previously reported estimates for *Tr*Cel7A ^13,14^.

**Figure 2:**
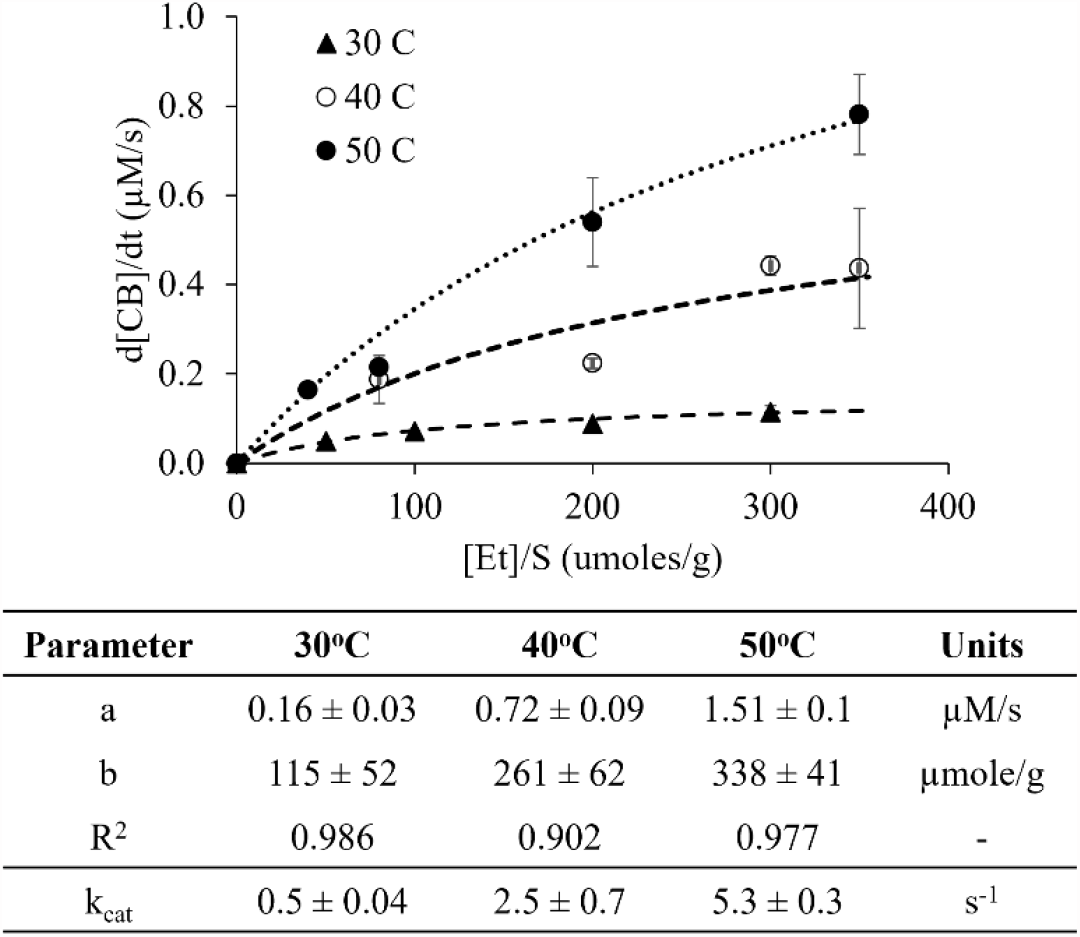
Initial rates of cellobiose release from swollen cellulose at varying TlCBHI loadings. Error bars reflect ranges for duplicate measurements. Lines fit to Equation 1 are shown, and fitting parameters given in the table. Standard errors of parameter estimates were calculated from 95% confidence intervals and propagated to estimates of k_cat_ calculated using Equation 2.

**Figure 3:**
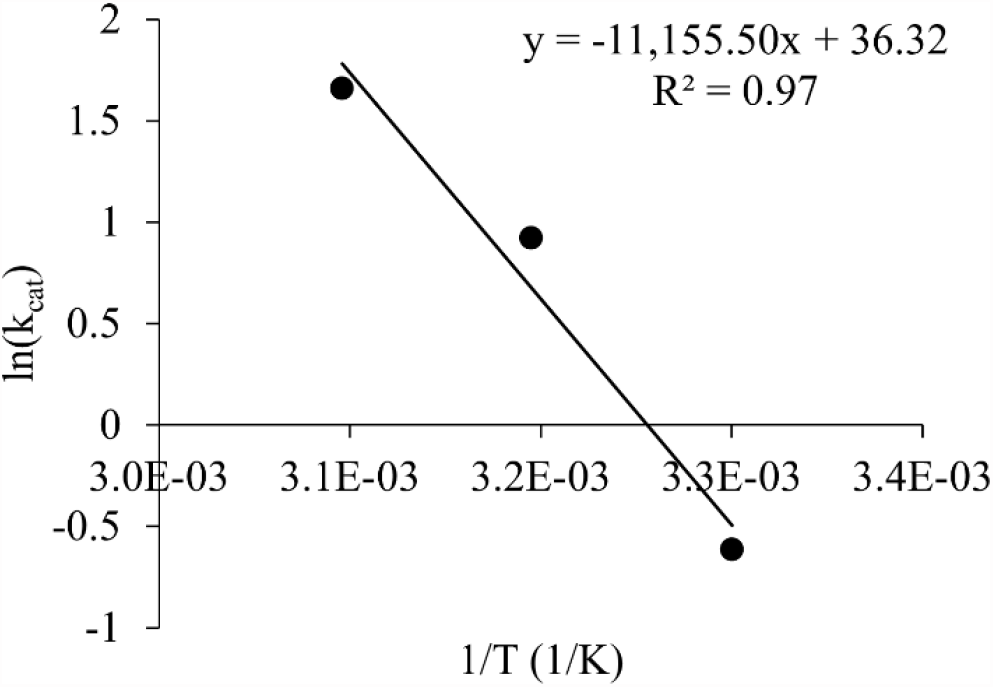
Double reciprocal Arrhenius plot used to estimate the energy of activation. The slope of the line fit was – 11,155.50 with a standard error of estimate of 2055.

## Conclusions

This short study characterized the catalytic nature of *Tl*CBHI, estimating that *Tl*CBHI hydrolyzes cellulose with a turnover rates (i.e. k_cat_) of *Tl*CBHI of 0.5 s^-1^, and 2.6 s^-1^, and 5.3 s^-1^ at 30 ^°^C, 40 ^°^C, and 50 ^°^C, respectively, with an energy of activation, E_a_, of ∼93 kJ/mol. Similar to the well characterized *Tr*Cel7A, *Tl*CBHI successfully complexed with and hydrolyzed easily accessible productive binding sites. However, *Tl*CBHI was either less stable for longer reactions than *Tr*Cel7A, or not as effective as *Tr*Cel7A at finding less accessible productive binding sites of cellulose.

## Acknowledgements

Jennifer Nill for purifying *Tr*Cel7A and preparing the swollen cellulose.

